# Local Mean Gradient Pattern (LMGP): A novel approach made for the classification of brain CT scan images

**DOI:** 10.1101/2024.08.06.606747

**Authors:** Kavya Singh, Anil Kumar Koundal, Navjeet Kaur

## Abstract

**Objective:** The visual descriptor methods like Local Binary Pattern (LBP) capture anatomical structures in captured images along with their disparities, which can be exploited by suitable methods for the diagnosis of medical anomalies. We developed a Local Mean Gradient Pattern (LMGP), based partly on LBP, a feature extraction algorithm for the classification of Computed Tomography (CT) images of the brain into normal, ischemic, or hemorrhage categories.

**Methods:** The ASID dataset comprises of 397 acute ischemic stroke CT scan images and the Kaggle dataset consists of 200 CT scan images, out of which, 100 images were of the abnormal brain type (hemorrhage), and the remaining 100 images which don’t show any abnormalities were categorized as normal cases. The results of LMGP were compared with eight other feature extraction methods using linear, sigmoid, polynomial, and RBF kernels of the SVM classifier.

**Results:** The effectiveness of our methodology was evaluated using recall, logarithmic loss, accuracy (ACC), confusion matrix and area under curve (ROC) metrics. The LMGP performed best when RBF-SVM was used for the classification and gave an accuracy of 93% and 95% in the case of five-fold and ten-fold cross-validation respectively. The ten-fold cross-validation gave a precision score of 0.9410, a recall score of 0.9521, an F1 score of 0.9458, and a negative log loss of -0.1645.

**Conclusion:** LMGP combines a unique and robust approach of CT scan image feature extraction by combining local information along with gradient change in pixels of the image. The present results of the study suggest the improved performance of the LMGP method over other methods compared in this study effectively.

## 1. Introduction

Brain hemorrhage or Intracranial hemorrhage refers to the accumulation of blood within the skull. It frequently results from trauma like motor vehicle accidents or other injuries, where blood vessels within the brain gets rupture. It may range from being relatively harmless to life-threatening, necessitating rapid surgical removal. A hemorrhage may potentially cause a stroke if it grows large enough to push against the brain [1]. Therefore, it is critical to find affected regions of the brain for timely therapeutic intervention. Medical imaging techniques including computed tomography (CT) and magnetic resonance imaging (MRI) are frequently used to diagnose anomalies of the brain, with CT scans usually being preferred for economic reasons.

A powerful tool with accurate predictive power is required for identifying brain abnormalities automatically. In this direction, numerous researchers are trying to develop tools and methodologies to accurately distinguish abnormal brain images from normal ones by utilizing advancements in image processing and machine learning techniques.

Computer-aided detection (CADe) refers to a class of computer systems that helps doctors and medical technicians in the interpretation of clinical images [2]. CADe can combine elements of medical image processing with computer vision and artificial intelligence to generate valuable support systems for the detection of medical conditions. For a particular medical condition, extracting meaningful information from medical images and combining them with a machine learning algorithm for their classification can generate good accuracy for its use in clinical practices. Several methods based on feature extraction that were proposed for the analysis of medical images have shown to be crucial for image segmentation and classification [3, 4]. Their relative ease and low computational requirements give them a substantial benefit over methods that are based on deep learning algorithms. These feature extraction methods can be based on statistical features, deep features, neighborhood relationships, colors, etc [2]. Local binary pattern (LBP), developed by Ojala et al., is one of the most widely employed methods for feature extraction purposes [5]. It has been widely utilized in classification problems because of its greater computational efficiency and indulgence with respect to the illumination changes. LBP was originally proposed for the texture analysis. Since it can only use a limited amount of structural knowledge, its performance is best when differentiating images of different textures and worst when distinguishing images of similar appearances (such as a tomato and an apple). It has been extensively used in a variety of applications, including biomedical image analysis, movement analysis, and image and video retrieval [6]. Furthermore, earlier approaches that have used texture features for the classification of images show a number of drawbacks, including indulgence with respect to the illumination variations and over-fitting as well as under-fitting issues with respect to the training and test datasets that can have a substantial impact on the classifier’s performances [4]. In this direction, we have proposed our methodology (LMGP as novel feature extraction method) for the classification of brain CT scan images into distinct categories and have compared the effectiveness of our findings to that of other eight feature extraction methods by using different classifiers.

## 2. Contribution of our proposed methodology

1. A new feature descriptor termed as local mean gradient pattern (LMGP) has been proposed for the classification of CT scan images in which the mean of the local image is computed and compared with the neighboring pixels of the central pixel value.
2. We used ASID and Kaggle datasets and tested our method on real patient data available in the form of CT scan images.
3. The efficacy of our proposed methodology (LMGP) was evaluated by comparing it with eight different feature extraction methods using different classifiers.

## 3. Related Work

Image texture delineates the pattern of spatial ordering of color or intensities in a given image or its selected region. A crucial component of computational image analysis is texture analysis, which allows for the efficient classification of images based on local spatial variations. Many different approaches to represent image texture have been put forth, with applications ranging from ground classification of aerial images to disease detection in biomedical images [7]. A common technique used for feature extraction of images is the local binary pattern (LBP) texture spectrum model proposed by Ojala et al. [5]. LBP is based on the relationship between the central pixel value with its neighborhood; if the neighbors have a value that is equal to or greater than the central pixel value, then their value is replaced by 1, and if they have a value that is less than the central pixel value, then neighboring pixel’s value is changed to zero. LBP generally operates by converting an image’s pixel value into a decimal number, often known as an LBP or LBP codes. It is employed not just in medical applications but also in computer vision and machine learning programs like facial image analysis, video and image retrieval, visual inspection, movement analysis, and others [5]. Apart from being less computationally intensive, it combines the structural and statistical properties of an image leading to an increase in texture analysis performance.

Researchers developed similar techniques after LBP, in order to address limitations in the original method and study the relationship between neighboring pixel values. Local ternary pattern (LTP) proposed by Tan et al. in which the neighboring pixel value is converted into a 3-value code (−1, 0, 1) instead of a 2-value code, which was in the case of LBP [8]. Center-symmetric local binary pattern (CS-LBP) proposed by Heikkil et al. also expanded LBP based on the consideration that the pixel values are compared with the opposing pixel values that are symmetrical with respect to that of the center pixel value [7]. Liao et al. presented dominant local binary patterns (D-LBP), an extended version of the LBP approach that efficiently captures the dominating pattern in texture images [9]. Images with complex geometries, such as corners and edges with extreme curvature, require the use of DLBP as it becomes unfeasible to classify these textures using uniform LBPs. Nanni et al. derived a variant of LBP by calculating the different shapes of the neighborhood using hyperbola, ellipse, circle, parabola and also considering binary as well as ternary encoding for the evaluation of the local-gray scale difference [10]. Guo et al. proposed a completed local binary pattern (CLBP) by combining CLBP_S, CLBP_M, and CLBP_C features into joint or hybrid distribution so that improvement can be made for texture classification [11]. Khellah et al. proposed a new variant for texture classification by merging global texture features with the local feature, which are obtained by using the LBP method [6]. Their fused texture features show a much higher rate of texture classification in comparison to the use of global or LBP features individually.

Unlike LBP, where each neighbor pixel value is converted to its binary form while comparing it with the central pixel which is further encoded to form local binary patterns, this methodology which is proposed by Zhao et al called Local Binary count (LBC) instead of encoding the binary values, the value of 1s is counted in the neighboring set [12]. Murala et al presented local ternary co-occurrence patterns (LTCoP), in which the appearance of similar ternary edges is calculated based on the comparison of central pixel value with respect to its surrounding neighboring set [13]. For the purpose of classifying textures, Zhang et al. suggested the local derivative pattern (LDP), which encodes directional pattern features based on local derivative changes using local second-order derivative features [14].

According to Murala et al. who proposed local mesh patterns (LMeP), each value of LMeP is calculated by comparing the surrounding neighbors for a given center pixel value in an image [3]. It is a brand-new feature extraction algorithm, inspired by the idea of LBP that is appropriate for indexing and retrieval of biomedical images. Local wavelet pattern (LWP), another feature descriptor derived by Dubey et al. computes the relationship between neighboring pixel values using local wavelet decomposition and ultimately considers its relationship with the center pixel value [4]. Furthermore, Dubey et al. presented the local diagonal extrema pattern (LDEP), a descriptor that allows for the calculation of the values and indexes of the local diagonal extremes in order to exploit the relationship among the diagonal neighbors [15].

A local binary pattern that is invariant to both rotation as well as scaling was proposed by Yuan et al. It included scale space, circular shift, and high-order directional derivative altogether. Various codes capitulate distinct orders, which eventually lead to the making of a distinct histogram for an image [16]. Another novel approach proposed by Tran et al. involves mainly two aspects which are features extraction followed by classification. Local binary pattern (LBP) and Local ternary pattern (LTP) are used for the purpose of feature extraction. Classification of features is done by a classification algorithm that functions on the basis of similarities among selected features. The Entropy-based Local Binary Pattern (ELBP) presented by Vidya et al. calculates the entropy of local regions within an image’s pixels before they are assembled into the Local Binary Pattern bin [17].

## 4. Background Knowledge

### 4.1 Local Binary Pattern (LBP)

LBP works by comparing the spatial relationship between neighboring sets of pixels with respect to the central pixel value of an image. LBP value is computed by comparing the central pixel value in an image with the neighboring sets of values (Figure 1). The neighboring value will be replaced by 1 if it is greater than or equal to the central pixel value, and it will become zero if it is less than the central pixel value. In figure 1, the method is shown with fictitious values.

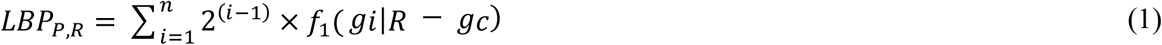

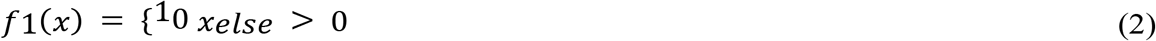

Center pixel of an image *I* has a grey value g_c_, g_p_ is the value of its neighbors and P is the total number of neighbors at a distance R from the central pixel (g_c_), where R is considered to be the radius of the neighborhood in an image. In figure 1, LBP code in a weights image (*I*_*w*_) of size 3×3 is calculated by 2^*i*−1^, where *i* refers to an index and ranges from 1 to 9 in a 3×3 matrix. Except for the center, all positions have weights set at the appropriate *Iw* positions. Lastly, the sum of the pixels is assigned to the center by multiplying the binary matrix with the weight matrix.

**Figure 1:**
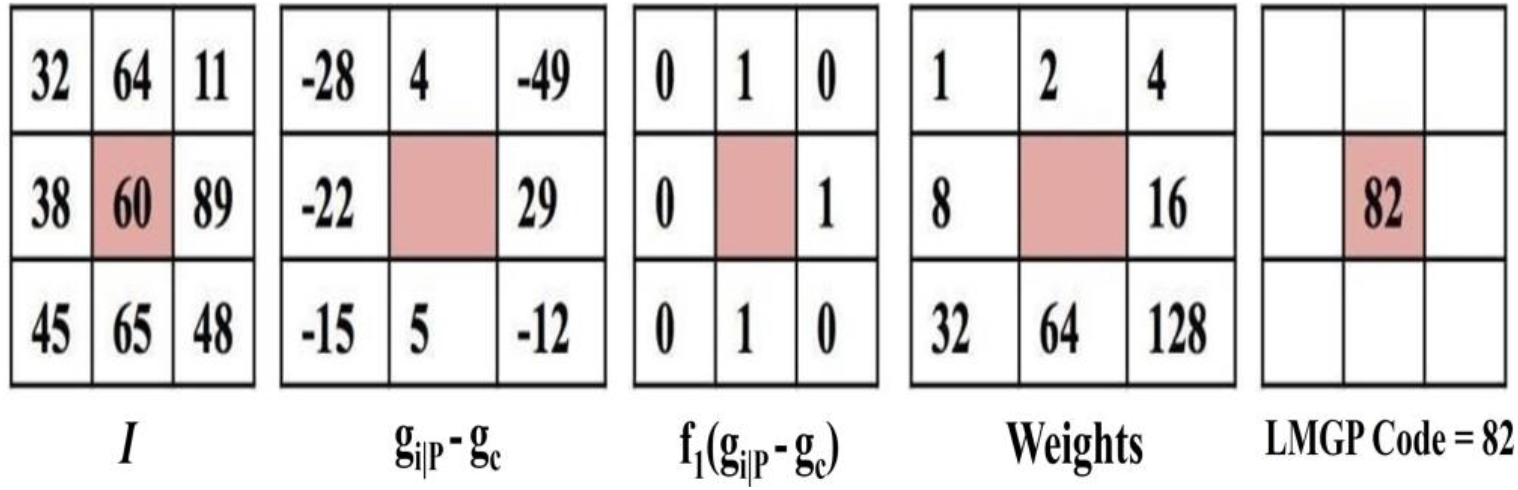
Representation of Local Binary pattern (LBP) for an image *I*.

## 5. Material and Methodology

### 5.1. Datasets

Kaggle (https://www.kaggle.com/code/anmspro/image-segmentation-brain-hemorrhage/input) dataset and AIS dataset (AISD) was used for the purpose of our study [18]. ASID dataset comprises of 397 acute ischemic stroke CT scan images that were obtained within 24 hours of the patient’s initial stage of symptoms. The ischemic stroke lesions present in the CT scan images were manually countered and traced by the doctors. In addition, the Kaggle dataset consists of 200 CT scan images, out of which, 100 images were of the abnormal brain type (hemorrhage), and the remaining 100 images which don’t show any abnormalities were categorized as normal cases. The slice thickness of CT scan images is 5mm. The size of the CT-scan images is 187 × 223 × 179 mm3.

### 5.2. Preprocessing

We have classified brain CT scan images based on the feature descriptor method LMGP. The steps have been described below:

Input data: CT scan images of acute ischemic stroke (AIS), hemorrhage, and normal patient. Result: prediction in the accuracy of classification model by using LMGP.

Pre-processing of CT scan images:

Due to the existence of numerous artifacts and disturbances, data obtained in the form of real-time images from the scan center are typically inappropriate for analysis. So, in order to improve the quality of an image, appropriate pre-processing techniques are required. It is considered as the preliminary step required for the image classification process. In our study, we have used the adaptive histogram equalization (AHE) technique, which can be applied to our patient data collected in the form of a CT scan image for improving the characteristics of an image. AHE is effective at improving local contrast and enhancing edges in CT scan images, which often have low contrast.

### 5.3. Proposed Methodology (LMGP) for feature extraction

This study proposes a unique feature extraction technique (LMGP) that considers the mean of a local image and the divergence of neighbouring pixels from the computed mean value. To the best of our literature survey done so far, this is the first attempt made for the classification of CT scan images.

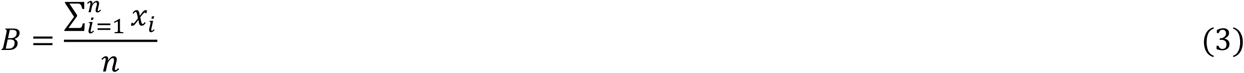

Where, B = mean of block

i = index value of block

n = total number of neighbours, xi = value of pixel at position i.

Gradient in x and y directions were calculated using following formulae:

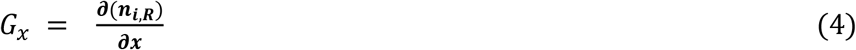

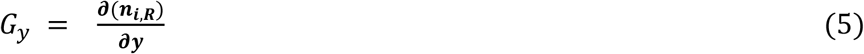

Gx = gradient in x direction

Gy = gradient in y direction

n_i,R_ = neighbour present at a distance R from the central pixel.

In a further step, we’ll compare the mean of local image with the gradient of all neighbours as shown in the equation below:

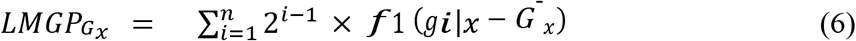

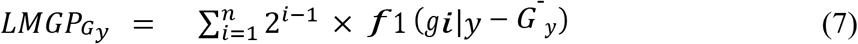

Where G^−^_x_ and G^−^_y_ are the mean of gradients in the x and y direction respectively.

g_i|x_ = gradient in x direction at position i.

g_i|y_ = gradient in y direction at position i.

f_1_ = function to calculate divergence of pixel values from average.

n = Total number of neighbouring pixels.

The equations 5 and 6 measure divergence of pixel gradients in X and Y direction respectively from the average gradient in each local segment of the image. The binary codes of all neighbours can be further reduced to LMGP code value by the equation given below:

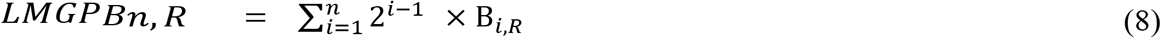

The equation represents the formula for calculating LMGP codes. *B*_*i,R*_ represents local image matrices obtained by thresholding after equations 6, and 7, and *i* is the index value of matrices. The equation converts threshold values to LMGP codes by assigning index values at a position which are above mean threshold values (Figure 2). These resulting code values are then added and assigned to the central pixel of the local partitions of the image. In the next step, all the calculated features, i.e. central pixel values are concatenated to form a single feature vector (*F*), inthe following order: *LMGP*_*B*_*n,R, LMGP*_*G*_*x*, and *LMGP*_*G*_*y*.

**Figure 2:**
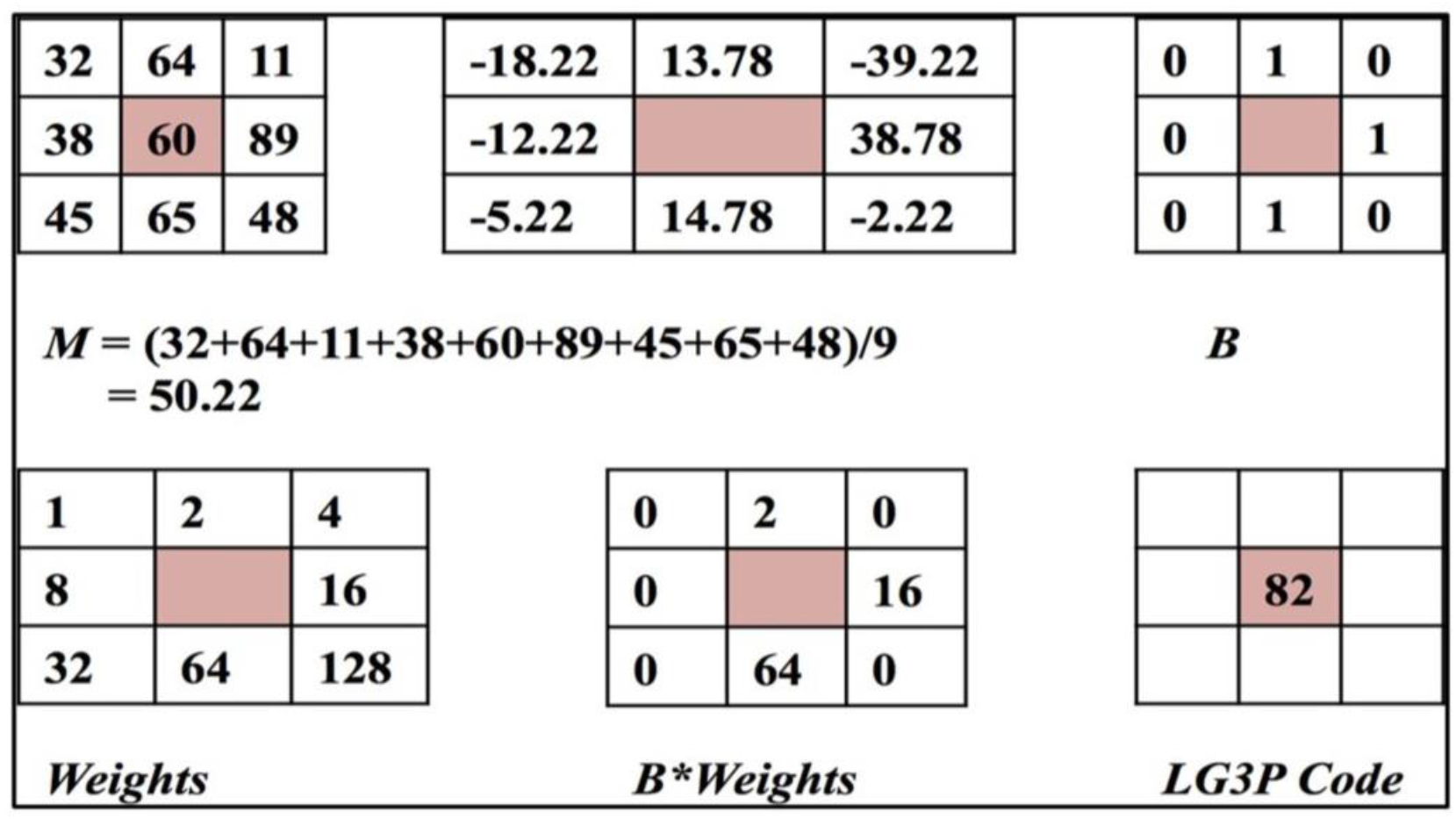
L.MGP codes of 3×3 size image.

In the final step, a discrete occurrence histogram is created from the calculated local code value for all pixels. The calculation of the histogram is given by the equation, which is described below:

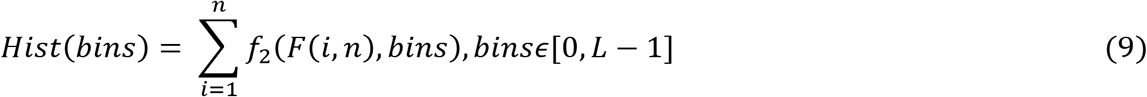

In our study we have taken bin size to be 256. This finally gives us a feature vector to be used as input for machine learning algorithms. In the case of LBP, its code value was calculated by comparing the central pixel with its neighbouring pixel values as shown in (Figure 1). For the calculation of LMGP, we have followed the same protocol where we have calculated the mean (M) of a local image, which is further compared with neighbouring pixel value as shown in (Figure 2). The numbers in the figure 1 and figure 2 are hypothetical, taken to explain LMGP methodology. The red color indicates the central value, and the other eight pixels are its neighbors. If the mean is less than the value of the neighbouring pixel, it will be considered as zero, and if it is greater than the neighborhood, it will be treated as one. The binary code obtained is multiplied by the weights assigned for each neighbouring pixel value and their summation gives the LMGP code value.

#### Feature extraction from pre-processed data

1. Compare all neighbors of central pixel with that of mean intensities so as to create the value of B as shown in equation 3.
2. In both x and y directions, calculate gradient *Gx* and *Gy* using Equations 4 and 5.
3. Calculate LMGP codes with Equations 6, 7, and 8.
4. Lastly, construct a histogram from the calculated code value using Equation 9.

### 5.4. Standard Scaling

Features were scaled by subtracting the mean and dividing the values by the standard deviation. Standard scaling is a popular choice for transforming data close to a normal distribution with a mean of around 0. Several machine learning algorithms like RBF-SVM, assume that all features are centered around 0.

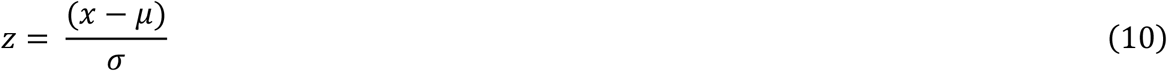

Equation 10 transforms extracted features by subtracting the mean from the value (*x*) and dividingoutcome quantity by *σ*, resulting in transformed feature values (*z*).

All the calculations were performed in Python and features were calculated using scikit-image library [19], and for training and testing datasets scikit-learn library [20] was used. Graphs were generated using matplotlib [21].

## 6. Results and Discussion

The proposed LMGP method is applied on the AISD and Kaggle datasets containing patients’ brain CT scan images [acute ischemic stroke, hemorrhage, and normal cases] (Figure 3) [18]. ASID dataset comprises of 397 acute ischemic stroke CT scan images that were obtained within 24 hours of the patient’s initial stage of symptoms. In addition, Kaggle dataset consists of 100 CT scan images of hemorrhage and 100 CT scan images of healthy patients. The slice thickness of CT scan images is 5mm. The size of the CT scan images is 187 × 223 × 179 mm3.

**Figure 3:**
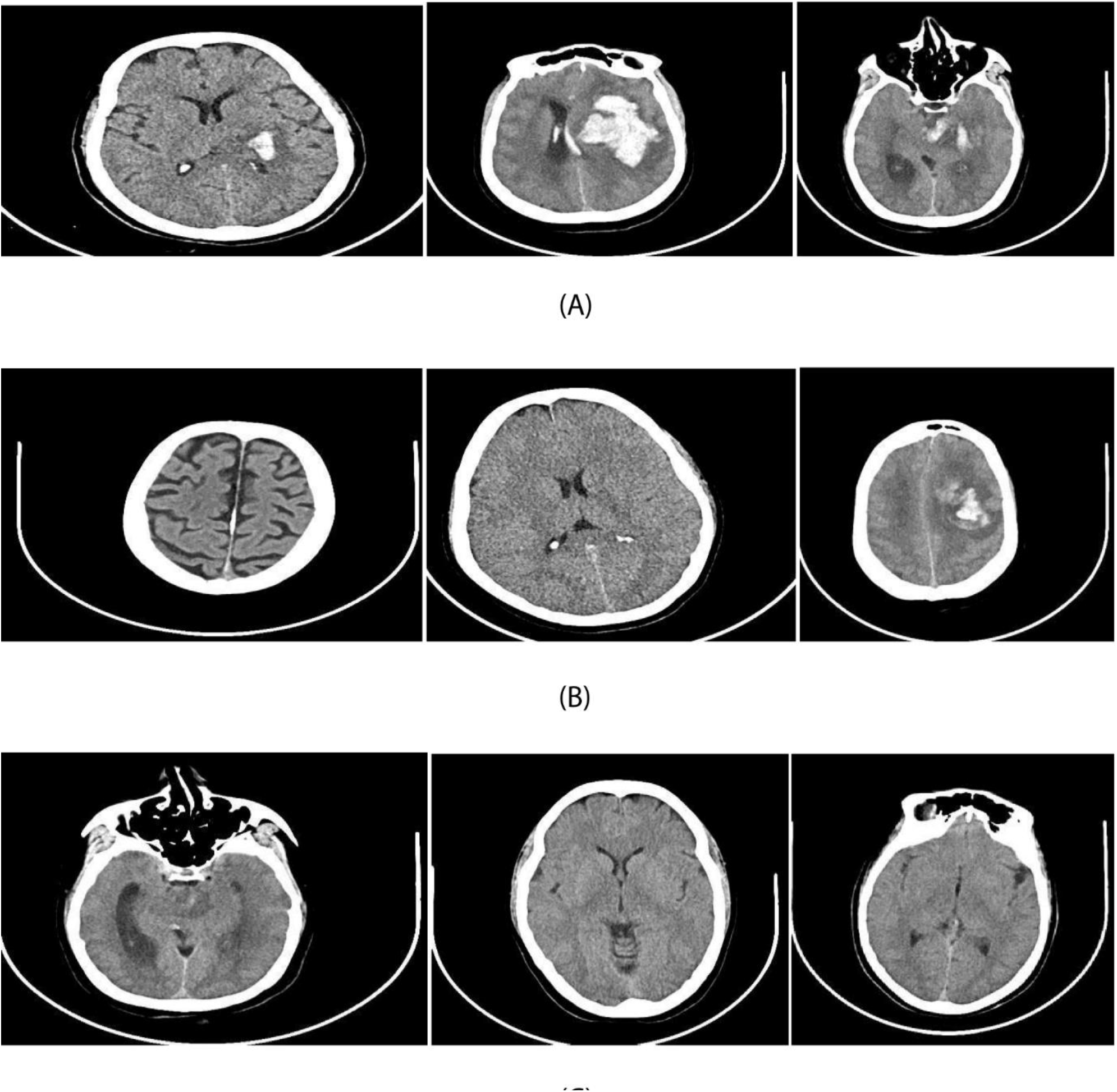
Sample images of (A) ischemic (B) hemorrhage and (C) normal cases.

In our study, 597 patients’ CT scan images were used for parameter tweaking and model training, and 104 CT scan images were used to test the proposed approach. Different classifiers were used for building the model. The effectiveness of our proposed methodology was evaluated on the basis of recall, logarithmic loss, accuracy (ACC), confusion matrix and area under curve (ROC) metrics. Before going into detail with the discussion related to the performance measure of different classifiers, let us look at some of the details related to the metrics that form the core in the process of validating our model. Logarithmic loss or simply log loss is often used as an evaluation metric. This basically works when the classifier assigns a probability to each class for the entire sample and has a log range (0, ∞). If the log loss is nearer to 0 it shows high accuracy whereas values further from zero indicate poorer accuracy of the model.

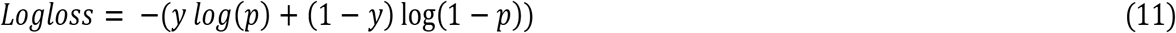

Where y represents sample i which can belong to either class 0 or 1 and p denotes the probability that sample y belongs to class 1.

Recall, also known as sensitivity, is defined as the division of the significant documents that are effectively retrieved.

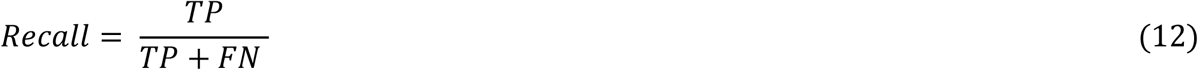

Area under the ROC curve (AUC) is a graph that is used to predict efficiencies of the classification model across all possible classification thresholds. This graph plots two parameters: True Positivity Rate (TPR) and Accuracy (ACC):

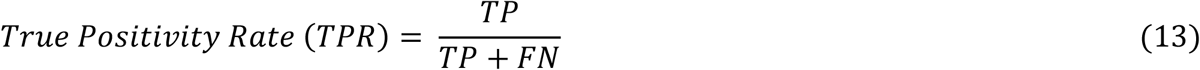

Accuracy (ACC) is used as another analysis metric for measuring the performance of classifiers for a particular feature extraction method.

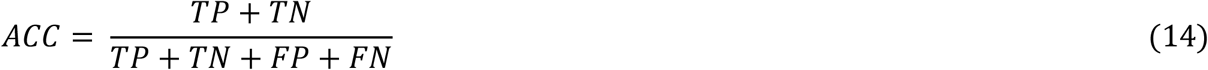

In equations 12, 13, and 14:

True positive (TP) is an output where the prediction of positive classes by the model is correct while false positive (FP) is an outcome in which positive classes prediction by the model becomes incorrect. True negative (TN) represents output in which the prediction of negative classes by the model is correct. In contrast to the above, false negative (FN) is an outcome where model prediction becomes incorrect in the case of negative classes.

In order to measure the performance of LMGP, several classification experiments using different classifiers were performed, i.e., linear SVM, sigmoid SVM, RBF SVM, and Poly SVM. The efficiency of LMGP is examined by comparing it with distinctive feature extraction methods, i.e., LTP, LWP, LBP, LDEP, LG2P, nLBP, TMEMPR, and GCLTP. For comparison, 5-fold and 10-fold cross-validation was performed on the dataset of CT scan images. In comparison to a simple leave-one-out cross-validation, the k-fold cross-validation provides a more precise estimation and variance of the accuracy. Overall, the LMGP performed best when RBF-SVM was used for the classification and gave an accuracy of 93% and 95% in the case of five-fold and ten-fold cross-validation respectively. The ten-fold cross-validation gave a precision score of 0.9410, a recall score of 0.9521, an F1 score of 0.9458, and a negative log loss of - 0.1645. The features were extracted using Python programming language using scikit-image library [19] and different variations of Support Vector Machine (SVM) were used for the classification using scikit-learn library [22]. The graphs were plotted using matplotlib [21].

### 6.1 Classification based on linear SVM

Linear SVM approach was used for the classification of brain CT scan images into two different categories, i.e., either abnormal (hemorrhage and ischemic) or normal patient cases. We have tested the accuracy of different feature extractors along with the LMGP method on five-fold and ten-fold cross-validation image datasets utilizing a linear SVM classifier. In five-fold cross-validation, the worst accuracy has been achieved by the LTP feature extractor, which is 71%, while others, apart from LMGP, have their accuracy between 79%-83% with LMGP having the highest at 86%. In the case of ten-fold cross-validation, again LTP performs the worst with an accuracy of 73%, precision score of 0.76 value, F1 score of 0.786, recall score of 0.81 value, and a least negative log loss of -0.605. So, in the case of both fivefold as well as tenfold, LMGP performs the best with an accuracy of 86% and 87% respectively. In addition to accuracy, other evaluating metrics like precision score, recall score, F1 score, and negative log loss also show that LMGP is better in terms of performance compared to other feature descriptors used for the classification of brain CT scan images based on Linear SVM classifier. Further information related to different metrics has been shown in table 1 and table 2. Table 1 and table 2 shows Accuracy, Precision, Recall, F1 Score, and Negative Log Loss obtained after validation using 5-fold cross validation and 10-fold cross validation respectively. The receiver operating characteristic (ROC) curve all feature extraction methods has been shown in figure 4 and figure 5, which correspond to 5-fold and 10-fold cross validation respectively.

**Table 1:**
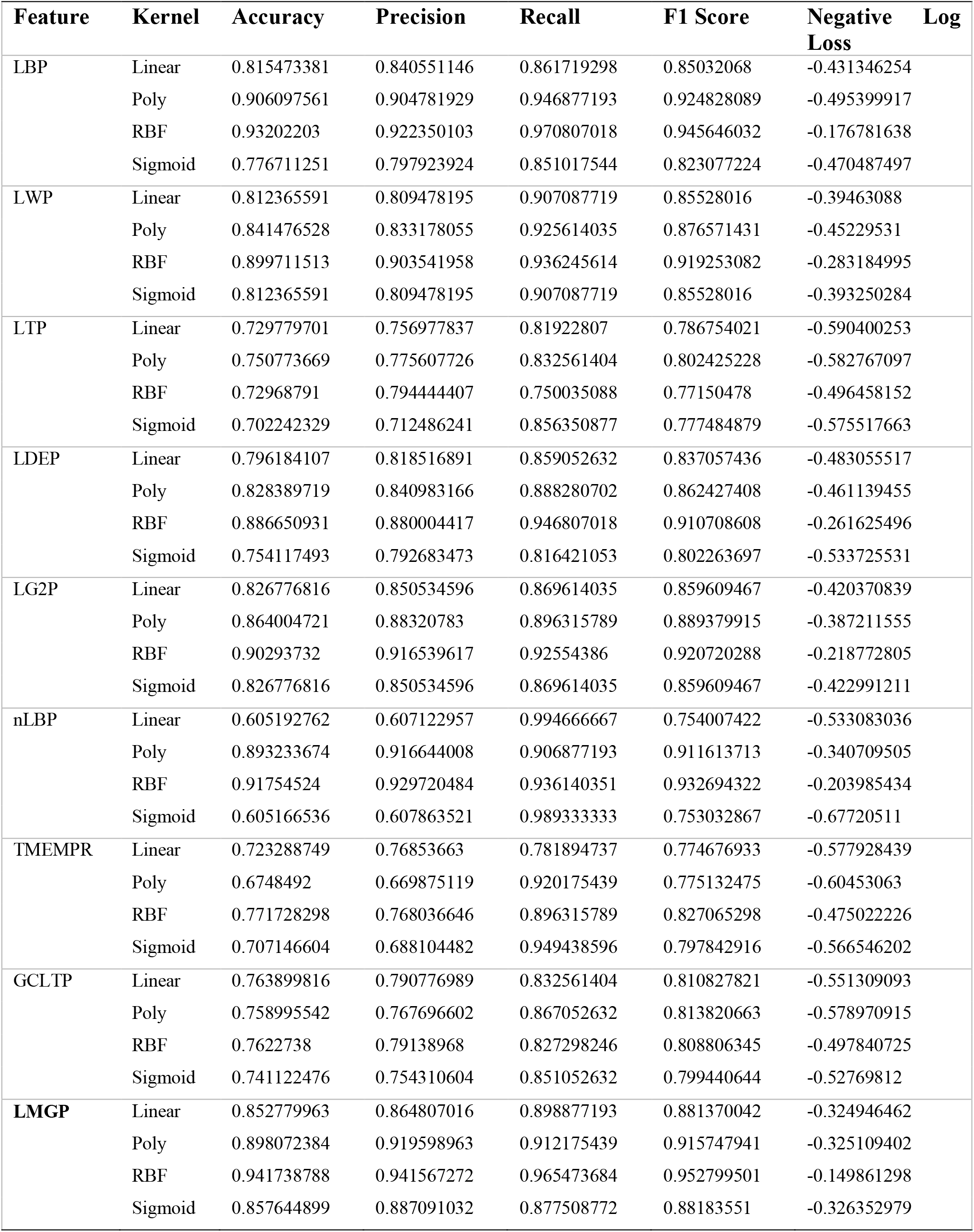
Metrics to evaluate our machine learning models in 5-fold cross-validation.

**Table 2:**
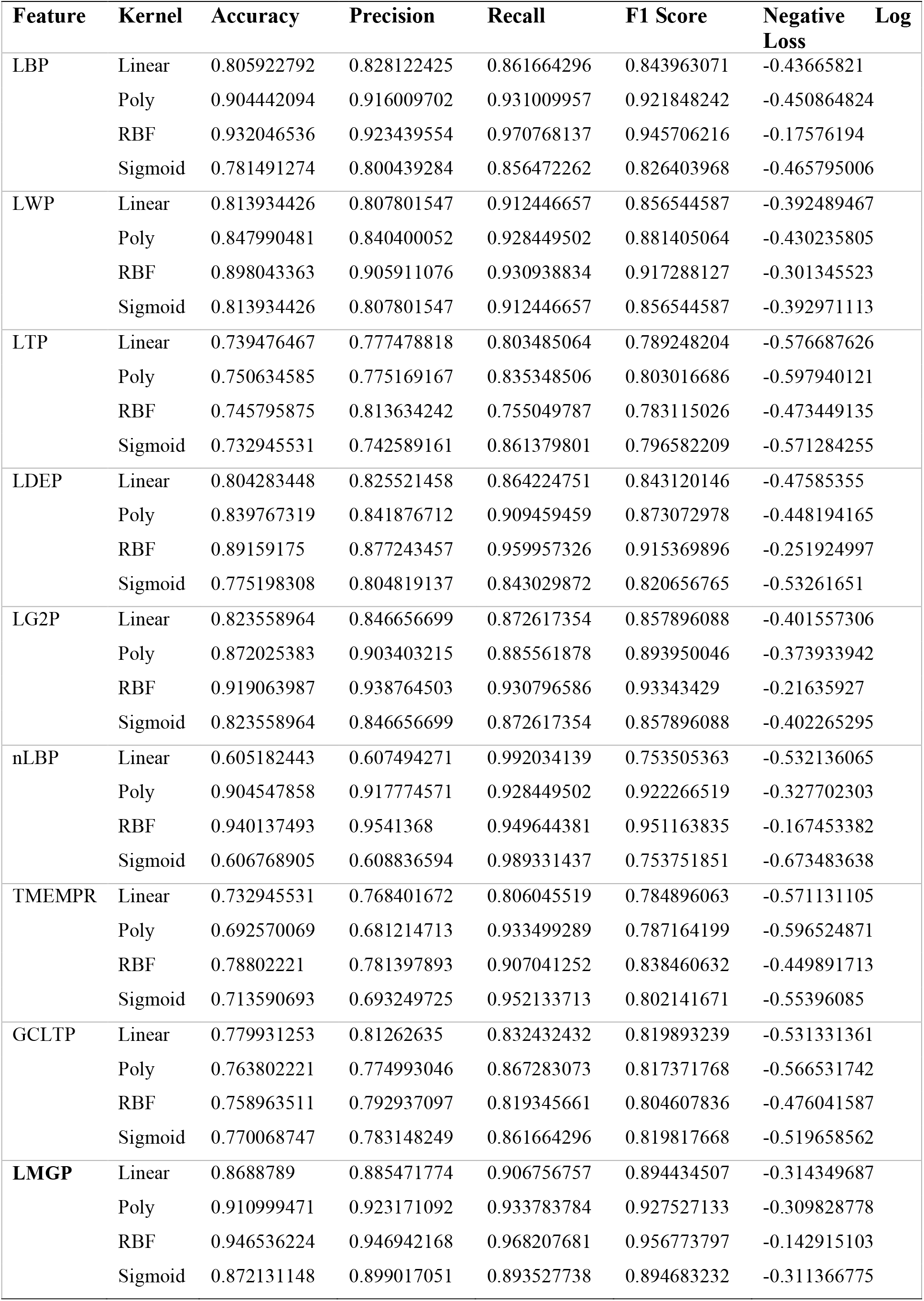
Metrics to evaluate our machine learning models using 10-fold cross-validation.

**Figure 4:**
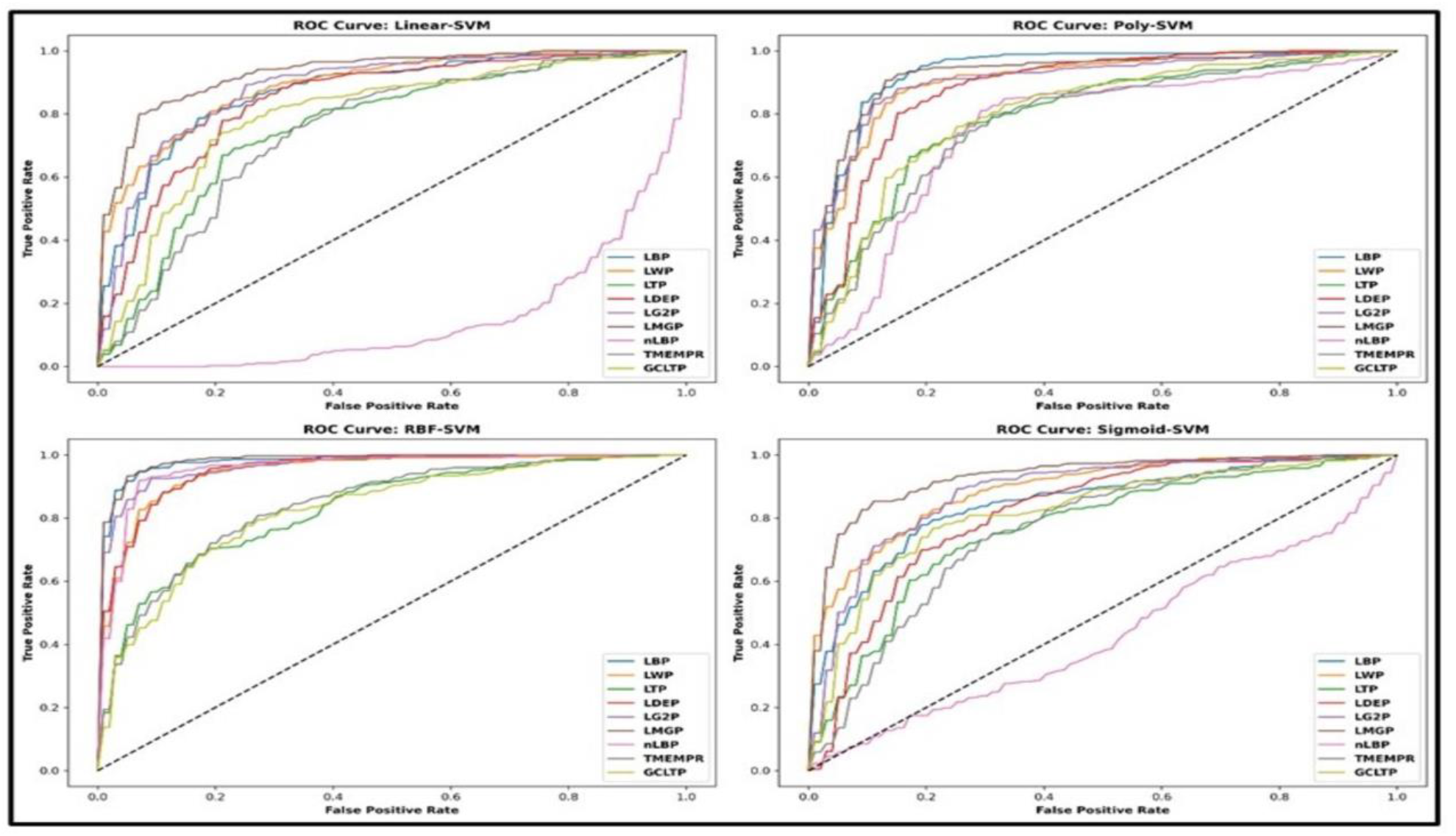
ROC AUC Curve of different SVM kernels using 5-fold cross-validation.

**Figure 5:**
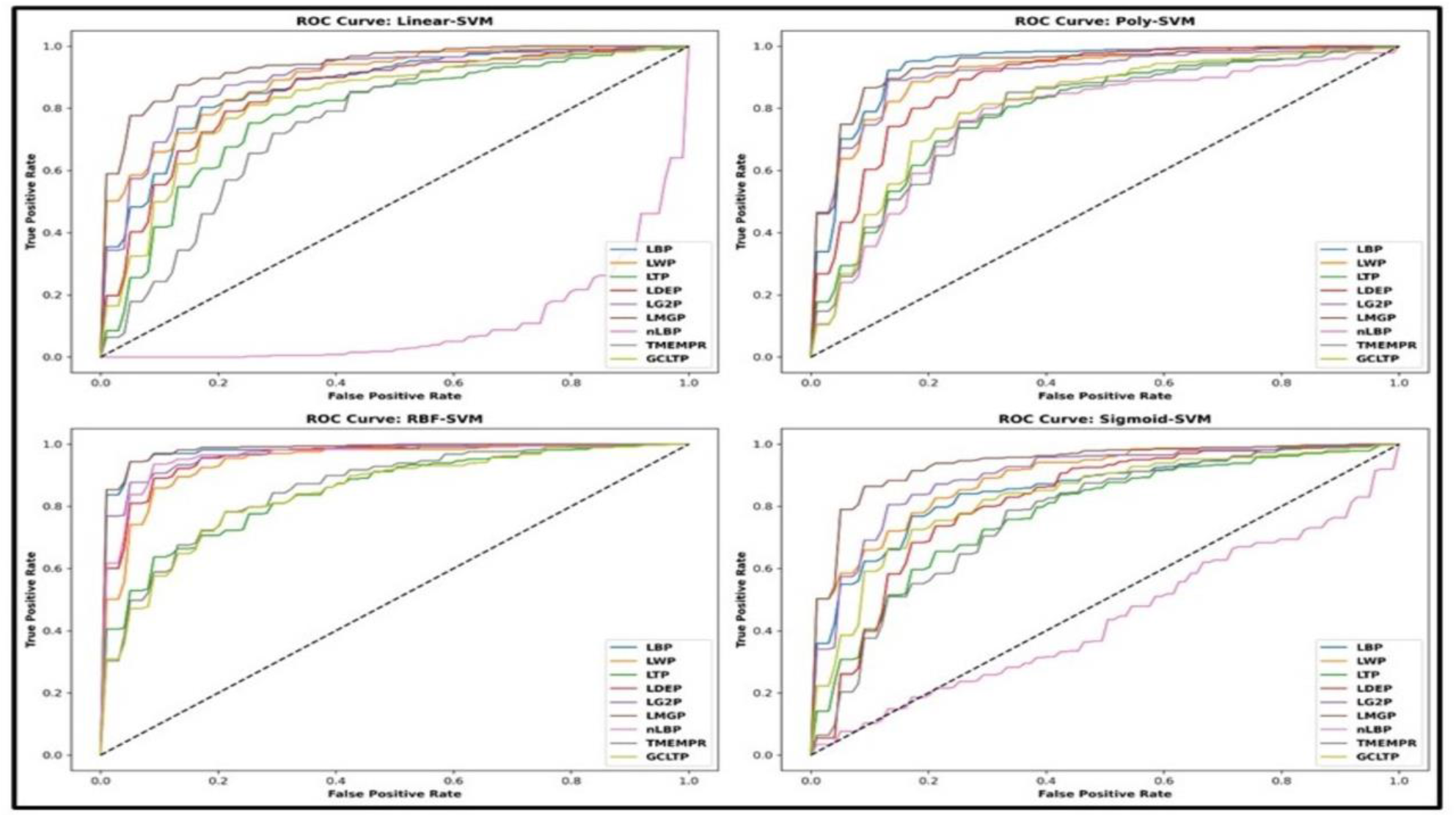
ROC AUC curve of different SVM kernels using 10-fold cross-validation.

### 6.2 Classification based on sigmoid SVM

The classification of Brain CT scan images by sigmoid SVM yielded 84% accuracy by LMGP, which was the highest observed in case of five-fold classification. LTP gave the worst accuracy of only 71%, while other feature descriptors give a performance with an accuracy that lies between 76% and 83%. In the case of ten-fold cross-validation, LMGP again gave the best accuracy of 87%, higher than other feature extraction methods, which had accuracies in the range of 73% to 83%. LMGP showed the best performance with an accuracy of 84% and 87% in the five-fold and ten-fold methods respectively. In the case of ten-fold cross-validation LMGP method gave precision score of 0.89 value, recall score of 0.92, F1 score of 0.89, and negative log loss of -0.31 which were slightly higher as compared to five-fold cross-validation results (Table 1 and Table 2). Hence, LMGP was the best feature extraction technique amongst methods in the study for classification of using the Sigmoid SVM classifier. The performance measures of different descriptors have been indicated in table 1 and table 2 and its ROC curve is illustrated in figure 4 and figure 5.

### 6.3 Classification based on polynomial SVM

In this methodology, the image classification of brain CT scan images was done using a polynomial SVM classifier. In both the cases of five-fold and ten-fold cross-validation, LTP feature descriptor performs worst in terms of accuracy with 74% and 75% respectively. LMGP performs best in the case of tenfold cross-validation with the highest accuracy of 91%, precision score of 0.9348, recall score of 0.9334, F1 score of 0.9334, and negative log loss of -0.2928. Whereas other feature descriptors have an accuracy that lies between 75% to 89%. LMGP shows the best performance for the classification of brain CT scan images (Refer to Table 1, Table 2, Figure 4 and Figure 5).

### 6.4 Classification based on radial basis function (RBF) SVM

In this experiment, image classification for the brain CT scan images was performed by employingradial basis function (RBF) SVM. The feature descriptor that performed worst in both cases of five-fold and ten-fold cross-validation was LTP with an accuracy of 75% and 74% respectively. In the case of five-fold cross-validation, LMGP classification accuracy is highest at 93% whereas, other feature descriptor’s accuracies lie between 75% to 88%. Similarly, in the case of ten-fold cross-validation, LMGP performs best with an accuracy of 95%, precision score of 0.9410, recall score of 0.9521, F1 score of 0.9458, and a negative log loss of -0.1645. Therefore, LMGP proves to be the best method suited for the classification of brain CT scan images using the RBF SVM classifier. Further information regarding the classification accuracy of different feature extractors in ROC curve has been shown in figure 4 and figure 5 and also in table 1 and table 2.

## 7. Conclusion

We have proposed a new feature descriptor method local mean gradient pattern (LMGP) for the classification of brain CT scan images. LMGP is an extension of the LBP feature extraction method. In our proposed methodology, the local image is segmented into the local blocks, and the mean of both gradients is taken and compared with the rest of the neighbouring pixel values of the block. In the next step, the binary code obtained is multiplied by the weights assigned for each neighbouring pixel and their summation gives the LMGP code value. Further, LMGP codes are calculated in order to get their histograms, which are finally concatenated with all the histograms to form a single vector of LMGP. Therefore, our methodology helps in the extraction of more pertinent information so as to enhance the classification process of brain CT scan images. The effectiveness of the proposed methodology was evaluated by using distinct classifiers and the outcomes were compared with different feature extraction methods. Conclusively, LMGP performs better as compared to the other methods used for the classification of brain CT scan images.

## Abbreviations

AIS: Acute Ischemic Stroke
AISD: Acute Ischemic Stroke Dataset
CAD: Computer Aided Detection
CLBP: Completed Local Binary Pattern
CS-LBP: Center-Symmetric Local Binary Pattern
CT: Computerized Tomography
D-LBP: Dominant Local Binary Patterns
ELBP: Entropy-based Local Binary Pattern
LBC: Local Binary Count
LBP: Local Binary Pattern
LDEP: Local Diagonal Extrema Pattern
LDP: Local Derivative Pattern
LMGP: Local Mean Gradient Pattern
LMeP: Local Mesh Patterns
LTCoP: Local Ternary Co-occurrence Patterns
LTP: Local Ternary Pattern
LWP: Local Wavelet Pattern
ML: Machine Learning
MRI: Magnetic Resonance Imaging
SVM: Support Vector Machine.

## Acknowledgments

K.S. is thankful to the Ministry of Human Resource Development (MHRD) fellowship provided during her PhD tenure.

## Author’s contribution

**KS**: Machine learning, Research design, Data collection, Writing manuscript. **AKK**: Machine learning, Manuscript editing. **NK:** Manuscript editing and Supervision.

## Data availability statement

Publicly accessible datasets were used in this study. This information can be found at GitHub (https://github.com/GriffinLiang/AISD) and Kaggle platform (https://www.kaggle.com/code/anmspro/image-segmentation-brain-hemorrhage/input).

## Ethical approval

Not applicable.

